# Neuromuscular dysfunction in patient-derived FUS^R244RR^-ALS iPSC model via axonal downregulation of neuromuscular junction proteins

**DOI:** 10.1101/2024.08.17.607965

**Authors:** Nicolai von Kuegelgen, Katarzyna Ludwik, Samantha Mendonsa, Christine Roemer, Erik Becher, Laura Breimann, Mara Strauch, Tommaso Mari, Sandrine Mongellaz, Binyamin Zuckerman, Nina Grexa, Anna Oliveras-Martinez, Andrew Woehler, Matthias Selbach, Vincenzo La Bella, Igor Ulitsky, Marina Chekulaeva

**Affiliations:** Non-coding RNAs and mechanisms of cytoplasmic gene regulation, Berlin Institute for Medical Systems Biology, Max Delbrück Center for Molecular Medicine, Berlin 10115, Germany; Department of Genetics, Harvard Medical School, Boston, MA, USA; Proteome Dynamics, Max Delbrück Center for Molecular Medicine, Berlin 13125, Germany; Departments of Biological Regulation and Molecular Neuroscience, Weizmann Institute of Science, Rehovot, Israel; BIMSB Light Microscopy platform, Berlin Institute for Medical Systems Biology, Max Delbrück Center for Molecular Medicine, Berlin 10115, Germany; Department of Experimental Biomedicine and Advanced Diagnostics, ALS Clinical Research Center, Laboratory of Neurochemistry, University of Palermo, Palermo, Italy

**Keywords:** Motor neuron, mRNA localization, RNA-binding proteins (RBPs), ALS

## Abstract

Amyotrophic lateral sclerosis (ALS) is a neurodegenerative condition characterized by the progressive degeneration of motor neurons, ultimately resulting in death due to respiratory failure. A common feature among ALS cases is the early loss of axons, pointing to defects in axonal transport and translation as initial disease indicators. Here, we established a FUS^R244RR^-ALS hiPSC-derived model that recapitulates the motor neuron survival and muscle contractility defects characteristic of ALS patients. Analysis of the protein and mRNA expression profiles in axonal and somatodendritic compartments of ALS-afflicted and isogenic control motor neurons revealed a selective downregulation of proteins essential for the neuromuscular junction function in FUS-ALS axons. Furthermore, analysis of FUS CLIP and RIP data showed that FUS binds mRNAs encoding these proteins. This work shed light on the pathogenic mechanisms of ALS and emphasized the importance of axonal gene expression analysis in elucidating the mechanisms of neurodegenerative disorders.

## INTRODUCTION

Neurodegeneration disorders are devastating diseases, predicted to become the second leading cause of death by 2040 due to population ageing (Gitler et al., 2017). Amyotrophic lateral sclerosis (ALS), also known as Lou Gehrig’s disease, arises from the progressive loss of motor neurons, leading to paralysis and respiratory failure, with no cure currently available. ALS cases of different etiology share defects in RNA metabolism, particularly in splicing and axonal mRNA transport (reviewed in Burk and Pasterkamp, 2019). Notably, mutations in the RNA-binding proteins (RBPs) TAR DNA-binding protein 43 (TDP43) and Fused in Sarcoma (FUS) are known causes of ALS, and pathologic TDP43 inclusions are found in the brain and spinal cord of 97% of ALS patients, suggesting that defects in RNA metabolism are a common feature in ALS. To date, most therapeutic approaches are aimed at protecting neuronal cell bodies from degeneration. However, in ALS axonal degeneration occurs prior to the death of neuronal cell bodies (soma) and correlates with the onset of functional decline (reviewed in Salvadores et al., 2017). The primary role of axonal degeneration in disease progression is supported by post-mortem studies (Fischer et al., 2004), mouse models (Frey et al., 2000) and electrophysiological observations (Dengler et al., 1990). Therefore, investigating local axonal gene expression in ALS is critical for future research.

Additionally, it is crucial to acknowledge the limitations of mouse models in neurodegenerative research. Although treatments in mouse ALS models have spurred over 50 clinical trials, none have translated into successful human therapies, pointing to fundamental differences in disease biology between humans and mice (Katyal and Govindarajan, 2017). Furthermore, mouse models fall short of representing sporadic ALS, which accounts for over 90% of cases (Patten et al., 2014). Hence, there is an imperative need to develop ALS models that reflect the genetic background of patients. Since the disease targets motor neurons, generating human neurons with the patient’s genetic background is feasible only through differentiation of patient-derived induced pluripotent stem cells (iPSCs) into motor neurons.

In this context, we tackled both aforementioned challenges. Using FUS^R244RR^-ALS patient fibroblasts, we established an iPSC line alongside an isogenic control where the FUS^R244RR^ mutation was corrected. These iPSC lines were then differentiated into spinal motor neurons through direct programming (Fernandopulle et al., 2018; Hester et al., 2011; Mazzoni et al., 2013; Shi et al., 2018; Son et al., 2011; Velasco et al., 2016). Our FUS^R244RR^-ALS model successfully replicated the motor neuron survival and muscle contractility impairments seen in ALS patients. Previously, we developed a technique for isolating neuronal subcellular compartments – specifically, cell bodies and neurites – for omics analyses (referred to as spatial omics, (Ciolli Mattioli et al., 2019; Loedige et al., 2023; Ludwik et al., 2019; Mendonsa et al., 2023; Zappulo et al., 2017). We adapted this approach to iPSC-derived motor neurons, separating their axonal and somatodendritic fractions for comprehensive omics analyses.

Analysis of the protein and mRNA expression levels in axonal and somatodendritic compartments of ALS and control motor neurons identified a preferential downregulation of proteins critical for neuromuscular junction functionality in FUS-ALS axons. Additionally, our examination of FUS CLIP and RIP datasets showed a marked enrichment of direct FUS targets among the downregulated proteins. This work illuminates the pathogenic mechanisms of ALS and emphasizes the crucial role of axonal gene expression analysis in understanding neurodegenerative disorders.

## RESULTS

### Efficient and rapid differentiation of human stem cells into motor neurons via direct reprogramming

To generate motor neurons from human stem cells, we relied on induced expression of transcription factors specific for motor neurons (MNs): NGN2, ISL1, and LHX3 (Fernandopulle et al., 2018; Hester et al., 2011; Mazzoni et al., 2013; Shi et al., 2018; Son et al., 2011; Velasco et al., 2016) (**Figure 1A**). The resulting motor neurons have been referred to as iMNs (induced MNs), iNIL (for induced <u>N</u>GN2, ISL1, and <u>L</u>HX3), or i^3^LMNs (for induced 3 factors lower motor neurons). In particular, we integrated a construct expressing these factors under the doxycycline-inducible promoter into the safe-harbor CLYBL locus (Fernandopulle et al., 2018) into human stem cell lines. As parental stem cell lines, we used human iPSC and embryonic stem cell lines (HuES-3 HB9::GFP, (Di Giorgio et al., 2008) to generate NIL-hiPSC and NIL-hESC lines, correspondingly (see Methods for details). In addition to the transcription factors, the integrated construct constitutively expresses the reverse tetracycline transactivator (rtTA) and a fluorescent marker mApple.

**Figure 1.**
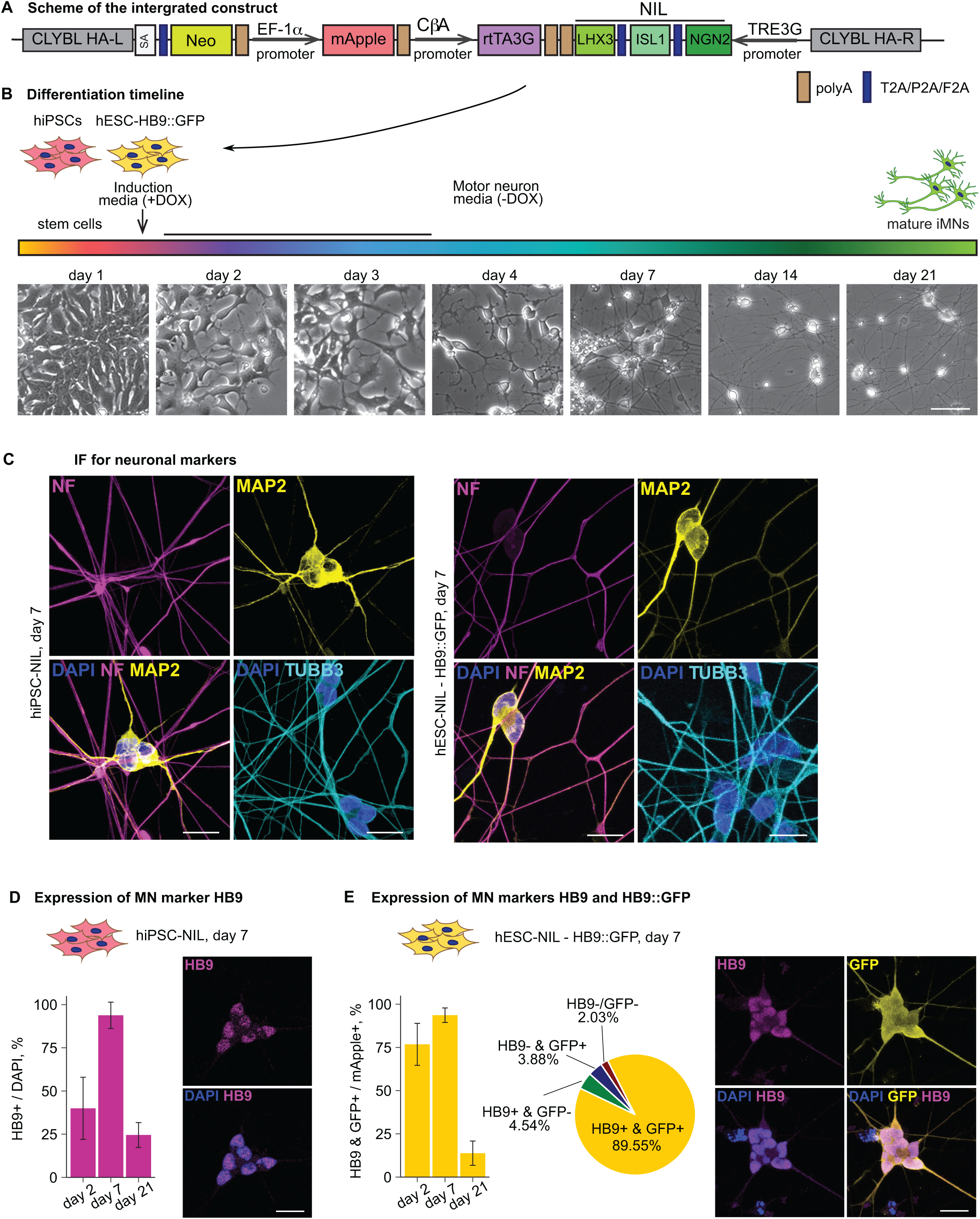
Direct programming is an efficient and robust protocol for the generation of motor neurons from hiPSCs and hESCs. (A) A scheme of the homology mediated repair template for insertion of NGN2-ISL1-LHX3 (NIL) cassette into CLYBL genetic locus. CbA: chicken beta-actin; SA: splice acceptor; HA L/R: homology arm left/right; Neo: neomycin resistance gene. (B) Differentiation protocol time course with exemplary images (Hues-3 line) illustrating morphological changes of the cells; DOX: doxycycline; iMN: human induced motor neuron. Scale bar = 100 µm. (C) iMNs express neuronal (TUBB3, cyan), dendritic (MAP2, yellow) and axonal (Neurofilament; NF, magenta) markers. iMNs derived from hiPSC-NIL (left) and hESC-NIL-HB9::GFP (right) on day 7 are shown. Scale bar = 20 µm. (D-E) Direct programming protocol produces a highly homogenous (around 90%) motor neuron culture. For the hiPSC-NIL differentiation, the percentage of HB9 positive cells and the exemplary immunofluorescence (IF) images at day 7 are shown in (D). For hESC-NIL differentiation, the percentages of HB9 and GFP positive cells and the exemplary IF images at day 7 are shown in (E). HB9: magenta, GFP: yellow, DAPI: blue, scale bar = 20 µm.

Induction of resulting NIL-hiPSC and NIL-hESC lines with doxycycline resulted in neuronal differentiation (**Figure 1B**). Cells started to change their morphology from day 2 (MN precursors), and long projections (neurites, or neuronal processes) extended from the cell body already by day 7 (early MNs). Neurites continued to elongate up to day 21 (mature MNs).

We confirmed the neuronal identity on day 7 of differentiation by immunostaining with neuronal (**Figure C**, TUBB3), dendritic (MAP2), and axonal (Neurofilament, or NF) markers. To validate the MN identity of resulting neurons and evaluate the efficiency of differentiation, we used immunostaining and imaging to analyze the expression of HB9, a transcription factor characteristic of early MNs (Arber et al., 1999). On day 7 more than 90% of cells were HB9-positive, both in the case of NIL-hiPSC-(**Figure 1D**) and NIL-hESC-derived neurons (**Figure 1E**). The NIL-hESC line also carries the green fluorescent protein (GFP) under the control of the *HB9* promoter: in these cells, GFP is only expressed when the *HB9* gene is active. Hence, GFP expression serves as an easily detectable visible marker for differentiation efficiency. Almost 90% of the NIL-hESC-derived neurons expressed both GFP and endogenous HB9 (**Figure 1E**). These data show that the expression of MN-specific transcription factors allows much higher differentiation efficiency (∼ 90%) than protocols based on small molecules (below 50%, Maury et al., 2015; Patani et al., 2011; Shimojo et al., 2015; Wichterle et al., 2002, reviewed in Patani, 2016).

### iMNs display expected differentiation patterns and are transcriptomically similar to primary mature motor neurons

To better characterize the differentiation process, we performed mass spectrometry and RNA sequencing (RNA-seq) analyses of different differentiation stages (**Figure 2**, **Table S1**). The proteomic and transcriptomic expression profiles of cell type markers, presented as heatmaps in **Figures 2A and B**, show a clear progression from stem cells via neuronal precursors to mature MNs (see **Figure S1** for global expression profiles).

**Figure 2.**
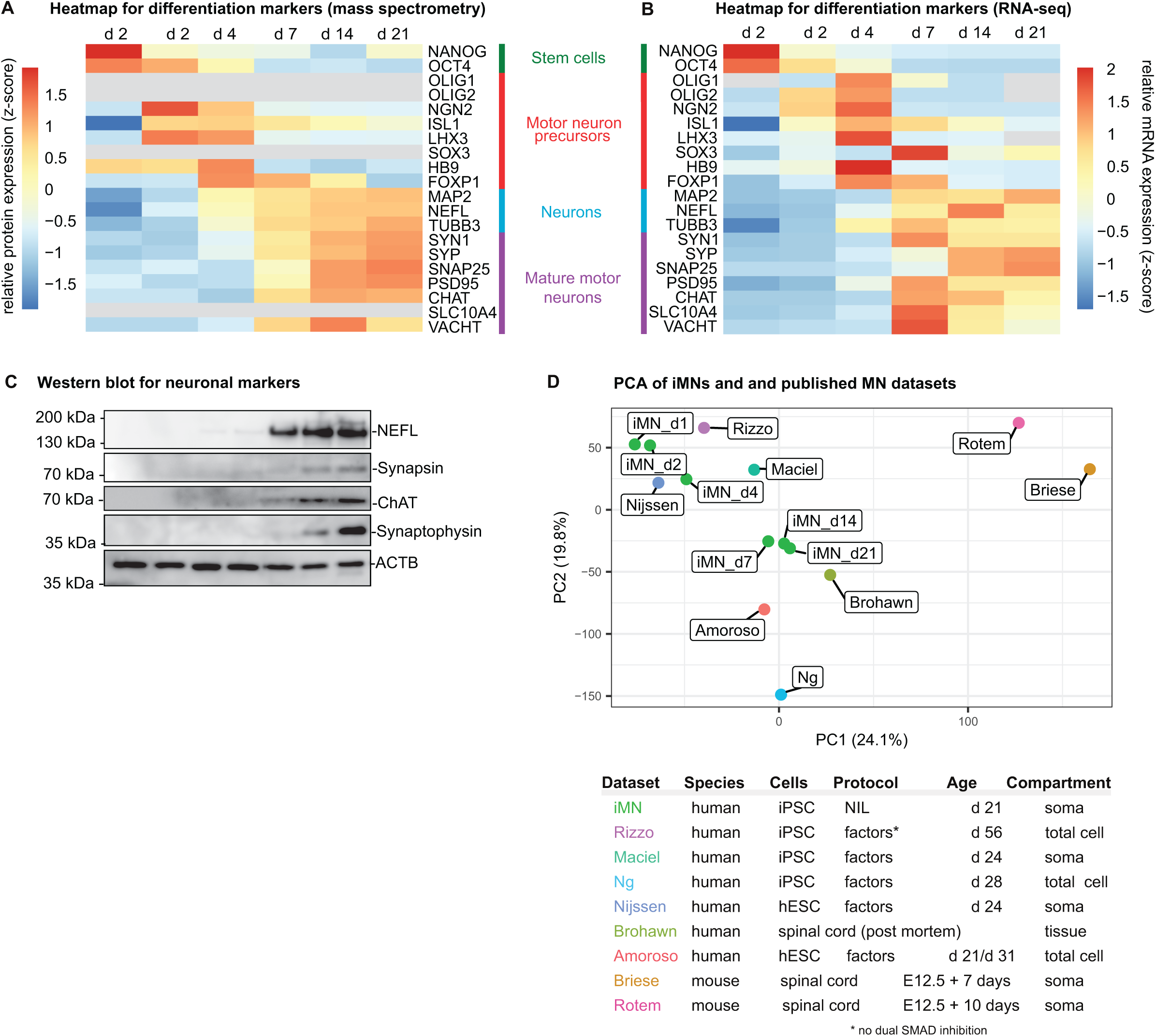
iMNs have MN marker signatures and are transcriptomically similar to primary MNs. (A-B) Heatmaps showing the expression of markers for stem cells (green), neuronal progenitors (red), neurons (blue), and mature motor neurons (magenta) along differentiation stages. Mass spectrometry values are shown in (A) and represent z-score transformed mean log2 LFQ (label-free quatification). mRNA-seq values are shown in (B) and represent z-score transformed mean log2 TPM (transcripts per million of library reads). (C) Western blot of mature neuronal (NF, Synapsin, Synaptophysin), and MN (ChAT) markers during differentiation. The position of size markers is indicated on the right. ACTB was used as a loading control. (D) iMNs are transcriptomically similar to primary cortical neurons. The principle component analysis of variance stabilization transformed (DESeq2::vsd) RNA-seq expression data from iMNs (this study) and publicly available datasets, presented in the table on the right. The table indicates the species, type of cells or tissues, age, subcellular compartment, and differentiation protocols used. “Factors” designates dual SMAD inhibition (unless marked*) and small molecule activation of the SHH pathway.

Stem cell markers (*NANOG* and *OCT4*) are well expressed in stem cells (**Figure 2A-B**, day 1) but were rapidly downregulated after doxycycline induction (days 2 and 4).

After doxycycline induction (day 2), MN-specific transcription factors *NGN2*, *ISL1*, and *LHX3* integrated into the NIL-hiPSC line become expressed (**Figure 2A-B**). These factors induce other neuronal precursor markers (*OLIG1*, *OLIG2*, *SOX3*, and MN-specific *HB9*) and quickly subside after doxycycline is removed at day 4. These data identify days 2-4 as the neuronal precursor stage. Expression of the HB9 protein was also validated by imaging (**Figure 1D-E**).

General neuronal markers MAP2, Neurofilament (NEFL) and TUBB3 reach high expression levels by day 7, corresponding to early MN stage. Finally, mature neuronal markers become highly expressed from day 7-14 of differentiation, reaching the highest level by day 21 (**Figure 2A-B**). Among those are components of synaptic vesicles: Synapsin 1 (SYN1), Synaptophysin (SYP), and Synaptosomal-associated protein 25 (SNAP25), as well as postsynaptic density protein 95 (PSD95) with a scaffolding role at the postsynaptic compartment. Moreover, at these stages the cells expressed known markers of mature MNs: Choline O-acetyltransferase (ChAT) (Oda, 1999) and Sodium/bile acid cotransporter 4 (SLC10A4). ChAT is the enzyme responsible for the biosynthesis of the neurotransmitter acetylcholine at the cholinergic synapses. SLC10A4 is involved in loading of synaptic vesicles and co-localizes with ChAT in in cholinergic neurons (Geyer et al., 2008; Larhammar et al., 2015). We validated the expression of mature MN and synaptic markers by western blotting (**Figure 2C**). Indeed, expression of synaptic (NEFL, SYN1, SYP) and mature MN markers (CHAT) reached the highest protein levels at day 21. Expression of these markers identifies the late stages of differentiation as mature motor neurons.

Next, we compared the transcriptome of our iMNs to the expression profiles of MNs in publically available transcriptomic datasets from primary and stem cell-derived MNs. For that, we used the same analysis pipeline (Wurmus et al., 2018) to process five RNA-seq datasets from human stem cell-derived MNs (Amoroso et al., 2013; Maciel et al., 2018; Ng et al., 2015; Nijssen et al., 2018; Rizzo et al., 2019), one dataset from human spinal cord (Brohawn et al., 2016) and two datasets from mouse spinal cord or primary motor neurons (Briese et al., 2016; Rotem et al., 2017). We compared the global similarities between the datasets using principle component analysis (PCA) (**Figure 2D**). Critically, our iMNs (day 7-21) clustered together with the human spinal cord samples (Brohawn et al., 2016). We detected more differences between spinal cords and MNs obtained with the differentiation protocols based on small molecules (Amoroso et al., 2013; Maciel et al., 2018; Ng et al., 2015; Nijssen et al., 2018; Rizzo et al., 2019). As expected, a clear separation between human and mouse data was observed in PCA. This analysis shows that our iMNs are transcriptomically similar to the mature human primary MNs and compare favorably to other available differentiation protocols in this regard.

### iMN are electrophysiologically active and form neuromuscular junctions

To ascertain the functionality of iMNs, we examined their expression profiles for the markers of electrophysiological activity. To achieve this, we used single-cell RNA sequencing data from patch-clamped neurons, which associated expression profiles with different action potential (AP) neuronal types (Bardy et al., 2016; Newman et al., 2019). In particular, some cells produced single spikes in patch clamping experiments (**Figure 3A**, left, low functionality: AP types 1, 2, 3, purple), other cells generated multiple spikes at low frequency (middle functionality: AP type 4, light-green), and there were also cells producing multiple spikes at high frequency (high functionality: AP type 5, dark-green). Higher neuronal activity correlated with upregulation of specific genes, for example, Piccolo (*PCLO*) required for neurotransmitter release at synapses (Wang et al., 2009), Trafficking protein particle complex subunit 6B (*TRAPPC6B*) involved in neuronal development (Marin-Valencia et al., 2018), and others (**Figure 3A**, left).

**Figure 3.**
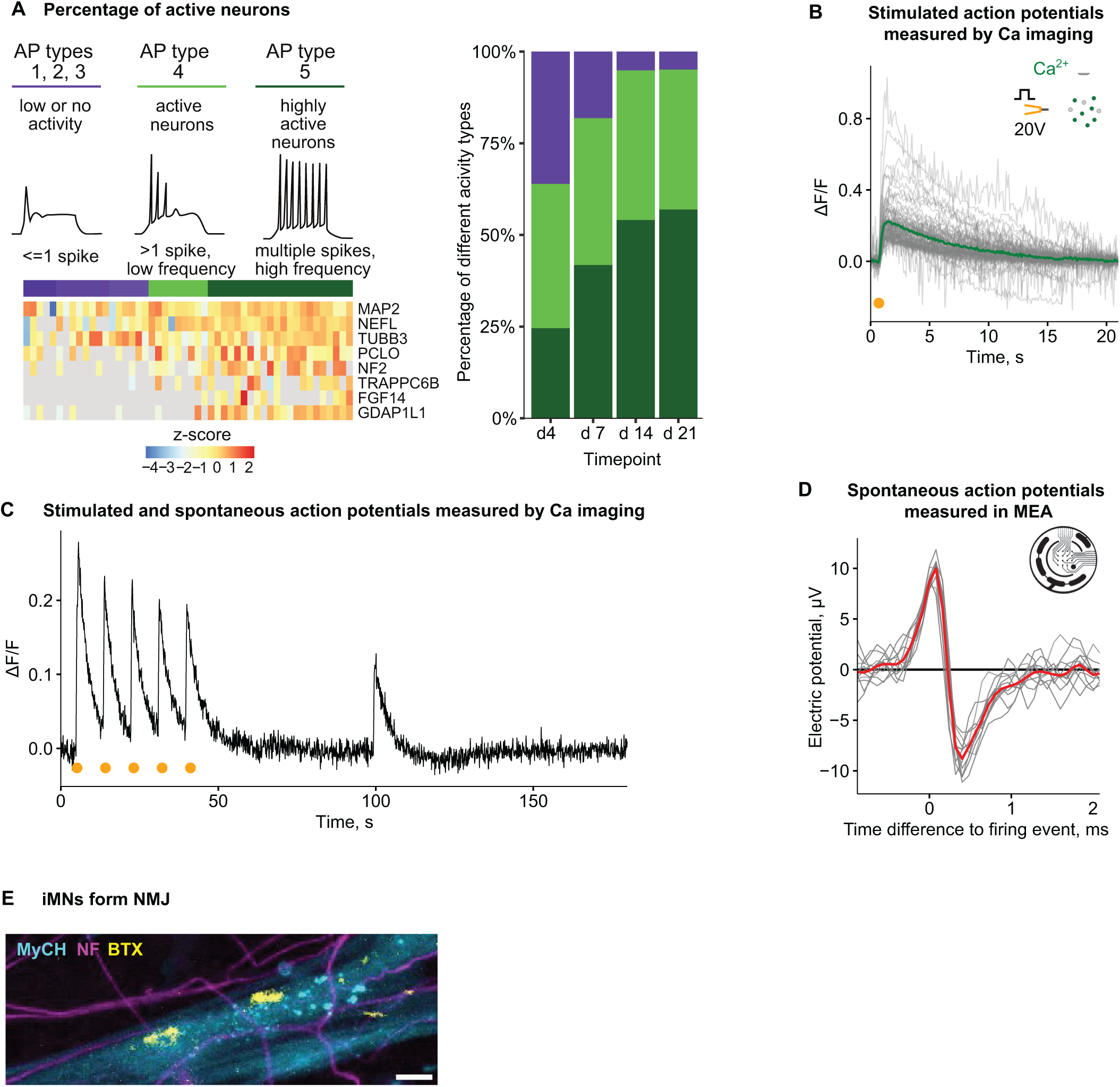
iMNs are electrophysiologically active. (A) Electrophysiological activity of iMNs, reflected in the frequency of spontaneous spikes, was predicted using scRNA-seq and electrophysiology data (Bardy et al., 2016). *Left:* Exemplary electrophysiology profiles of cells corresponding to different functional types (adapted from Bardy et al., 2016) and heatmap showing z-score transformed expression values of general neuronal (MAP2, NEFL, TUBB3), synaptic (PCLO, NF2) and active transmission markers (TRAPPC6B, FGF14, GDAP1L1) used to determine the functional type. *Right:* Bar plot showing percentages of cells matched to each functional type for different differentiation stages. (B-C) Neuronal activity of iMNs recording with calcium imaging. (B) Calcium transients after induction of activity potentials in iMNs (day 20 after the start of differentiation) using 20 V stimulation. Fluorescence intensity changes from CalBryte520 are normalised to pre-stimulus baseline level, and individual measurements from 84 ROIs (grey) as well as the average (green) are shown. The inset shows a graphical representation of the experimental setup. (C) Exemplary calcium transient of a single region of interest (ROI) over the 1 min induction and 2 min observation period. 20V stimulation occurred at 5, 14, 23, 32 & 41 s time points (yellow marks), and changes in fluorescence intensity, normalized to baseline level, are shown. (D) Action potentials measured in iMN grown in an Axion multiwell microelectrode array (MEA) plate. Multiple spikes from the same electrode as well as the average (red, n = 20) are shown. The inset shows a graphical representation of the MEA plate. (E) iMNs form neuromuscular junctions in co-cultures with myotubes. Axons are visualized with NF staining (magenta), myotubes – with α-myosin heavy chain (MyHC) staining (cyan), and NMJs -with BTX staining (yellow). Scale bar = 10 μm.

Relying on these data, we applied cell-type deconvolution (Newman et al., 2019) to our RNA-seq data produced from different differentiation stages of iMNs. This analysis showed that the functionality of iMNs increases as cells proceed through differentiation, with more than 90% of functional neurons (light-and dark-green bars) at day 21 (**Figure 3A**, right).

We then experimentally measured the electrophysiological activity of iMNs using calcium imaging (**Figure 3B**). Electrophysiological activity in neurons is accompanied by an influx of calcium ions, which can be measured with calcium indicators -fluorescent molecules that respond to the binding of calcium ions. Using electrical stimulation, we could evoke action potentials in mature iMNs, visualized with calcium transients after stimulation (**Figure 3B**). Moreover, 67% of cells responding to electrical stimulation showed spontaneous activity potentials (**Figure 3C**). In addition to calcium imaging, we used a multielectrode array (MEA, Axion) to measure the electric activity of mature iMNs (**Figure 3D**). These experiments confirmed that iMNs generate spontaneous action potentials with typical de-and repolarisation patterns.

Another hallmark of functional mature MNs is the ability to form neuromuscular junctions (NMJs), specialized synapses between motor axons and muscle cells. To test if iMNs can form NMJs, we co-cultured them with muscle cells generated through the inducible expression of the myogenic transcription factor MyoD (Davis et al., 1987) using available protocols (Uchimura et al., 2017). The functionality of these neuromuscular cultures has been confirmed by evidence of muscle contractions and their responsiveness to curare (**Supplementary movie 1**), a drug that affects the interaction between motor axons and muscles (Bowman, 2006). To visualize NMJs, we stained the co-cultures for nicotinic acetylcholine receptors (nAchR), which serve as a marker of NMJ. nAchR is recognized with a fluorophore-conjugated bungarotoxin (BTX, **Figure 3E**). Indeed, we observed the formation of NMJs (BTX, yellow) at the sites of contact between motor axons (NF, magenta) and myotubes (myosin heavy chain, or MyCH, cyan). Thus, our analysis points to the hallmarks of functionality of iMNs – they exhibit electrophysiological activity and form functional NMJs.

### Efficient enrichment of motor axons for omics analyses

iMNs represent a promising cellular model to study motor neuron diseases, such as Amyotrophic Lateral Sclerosis (ALS), Spinal Muscular Atrophy (SMA), Charcot-Marie-Tooth disease (CMT), and others. Although it has been known for some time that the first signs of neurodegeneration in motor neuron diseases manifest in axons (Arendt, 2009; Dengler et al., 1990; Fischer et al., 2004; Frey et al., 2000; Selkoe, 2002), most prior studies in the field focused on whole neurons, therefore failing to detect local axonal changes. Therefore, we decided to adopt the spatial transcriptome and proteome analysis we developed earlier (Ciolli Mattioli et al., 2019; Loedige et al., 2023; Ludwik et al., 2019; Mendonsa et al., 2023; Zappulo et al., 2017) to iMNs.

To separate iMNs into axons and soma, we grew them on a microporous membrane so that soma stayed on one surface, and axons extended through the pores on another membrane surface (**Figure 4A**). We confirmed efficient separation by immunostaining of MNs growing on the membrane (**Figure 4B**). Importantly, staining with axonal (NF) and dendritic (MAP2) markers confirmed that with this approach we can isolate motor axons, primarily affected in motor neuron disease, rather than a mixture of axons and dendrites. We also validated the enrichment of selected axonal (NF, GAP43) and somatic markers (Histone H3, RNA II polymerase POLR2A) by western blotting on isolated subcellular compartments (**Figure 4C**).

**Figure 4.**
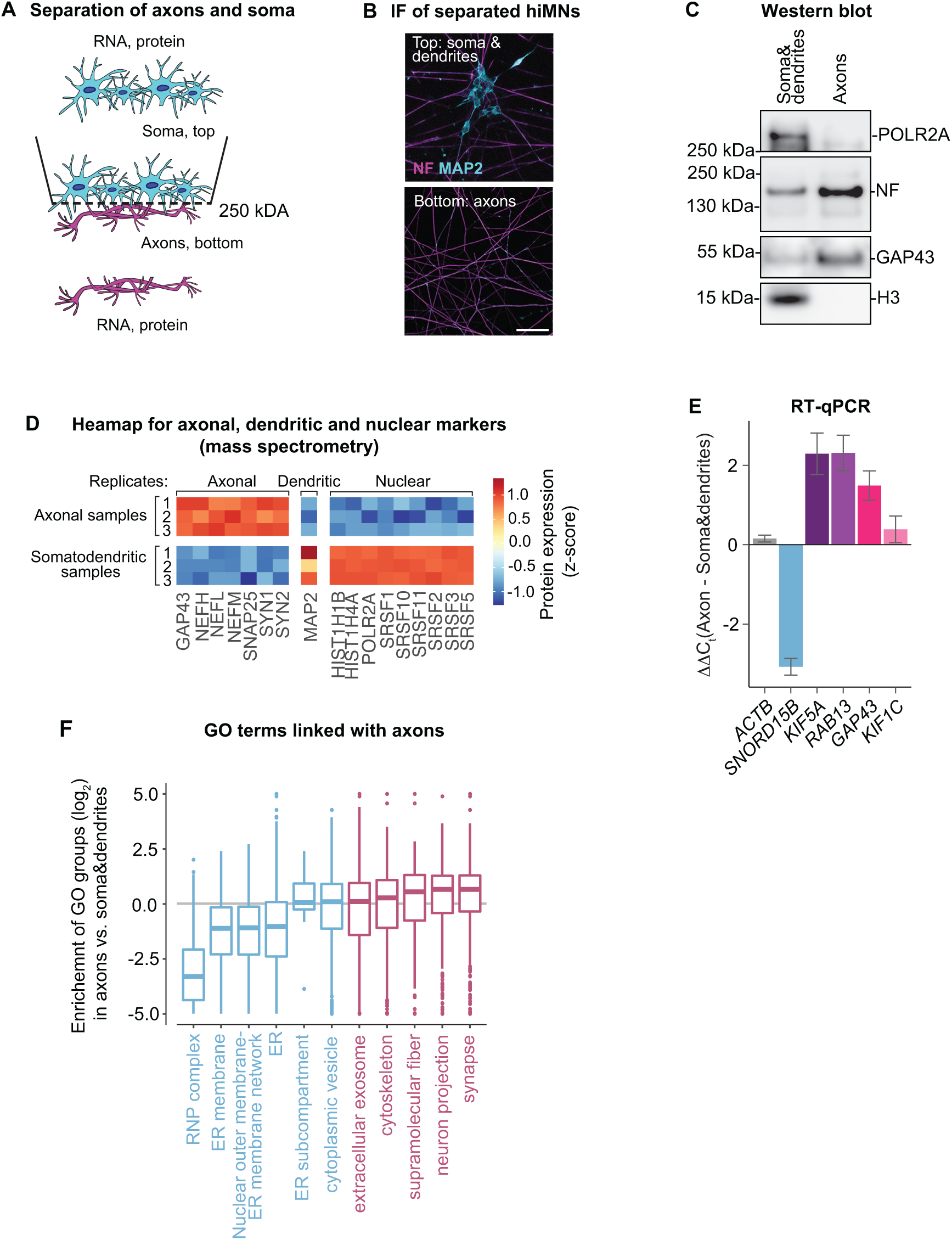
The local transcriptome and proteome of motor axons are linked with synaptic functions. (A) Scheme showing separation of iMNs on subcellular compartments. iMNs are grown on a microporous membrane with cell bodies (Soma) restricted to the top and axons extending to the bottom, allowing for the separation of the two compartments. (B) Exemplary image of iMNs grown on a microporous membrane: dendrites and soma (MAP2, cyan) are restricted to the top of the filter, while axons (NF, magenta) extend to the bottom. (C) Western blot analysis validating enrichment of somatic and axonal proteins in the separated compartments. Axons are enriched in neurofilament (NF) and GAP43 and depleted of histone H3 and RNA polymerase II (POLR2A). Position and size markers are indicated on the left. (D) Heatmap showing expression of axonal, dendritic, and nuclear markers in three biological replicates of axonal and somatodendritic samples. Plotted and z-score transformed label-free quantification (LFQ) values obtained by mass spectrometry analysis. (E) qRT-PCR analysis validates the enrichment of somatic and axonal RNAs in separated subcellular compartments. Nuclear RNA SNORD15B is enriched in soma, while ACTB mRNA is equally distributed, and RAB13, GAP43, KIF1C and KIF5C show different degrees of enrichment in axons. The difference in expression levels (ΔΔCt) of indicated transcripts between axonal and somatodendritic compartments samples is plotted on (Y). 18S rRNA was used as a normalization control. Bar, mean +/-SD, n = 3. (F) GO analysis indicates the enrichment of proteins with nuclear function in the soma and synaptic or neuronal functions in axons. Boxplots show the protein localization values (Axon/Soma log2 fold change) of all proteins belonging to selected GO terms. Terms were selected from GO cellular compartments if they showed a significant overrepresentation of proteins or genes localized to either compartment in both the proteomic and transcriptomic data.

We then proceeded to isolate axons and somatodendritic compartments from either side of the membrane. To identify their local proteomes and transcriptomes, we performed liquid chromatography-tandem mass spectrometry (LC-MS/MS) and RNA-seq of isolated subcellular compartments. We measured 5418 proteins using a label-free quantification (LFQ) method (Cox et al., 2014) and 15610 transcripts using RNA-seq (**Tables S2-S3**, see **Figure S2A-D** for quality controls). We performed differential expression analysis to identify proteins and transcripts that are differentially localized between two subcellular compartments. We defined axon-and soma-localized proteins and mRNAs based on their fold enrichment in axons versus somatodendritic compartment: axon-localized proteins and mRNAs are defined as significantly enriched in axons by at least 2-fold (**Figure S2E-F**, magenta), and soma-localized mRNAs (cyan)-as enriched in somatodendritic compartment according to the same criterion. Importantly, analysis of three biological replicates of isolated compartments confirmed consistent enrichment of axonal markers in axonal samples and dendritic (MAP2) and nuclear markers – in somatodentritic samples (**Figure 4D**), additionally validating our ability to enrich the axonal fraction. Among axonal markers were neurofilaments (NEFL, NEFM, NEFH), a major component of axonal growth cones Neuromodulin (Growth-associated protein 43, or GAP43 (Goslin et al., 1988)), components of synaptic vesicles synapsins 1-3 (SYN1, SYN2, SYN3) and SNAP25 (Suh et al., 2010). Nuclear markers included histones (HIST1H1B, HIST1H3A, HIST1H4A), a subunit of RNAII polymerase (POLR2A), and splicing factors (SRSF1, SRSF2, SRSF3, SRSF5, SRSF10, SRSF11).

Additionally, at the mRNA level, transcripts encoding several axonal markers were enriched in axons (**Figure S2F**, magenta). These included *Gap43* (Goslin et al., 1988), kinesin motor proteins involved in axonal transport such as kinesin-like protein KIF1C (*Kif1c*, (Dorner et al., 1998) and kinesin heavy chain isoform 5A (*Kif5a*, (Wang and Brown, 2010), which has previously been reported to be axonally enriched (Gumy et al., 2011) and a major regulator of neurodegeneration (Shah et al., 2022); *Rab13*, encoding a component of transport vesicles co-localizing with GAP43 (Di Giovanni et al., 2005), enriched in neurites across multiple types of neurons (Costa et al., 2020; von Kugelgen and Chekulaeva, 2020). We validated the axonal enrichment of these transcripts by RT-qPCR on RNA isolated from motor axons and soma (**Figure 4E**, purple to red bars). In contrast, nuclear marker small nucleolar RNA 15b (*Snord15b,* blue bar) was somatically enriched in our RT-PCR analysis.

To functionally characterize local proteomes and transcriptomes of iMNs we performed Gene Ontology (GO) overrepresentation analysis on proteins and transcripts enriched in one of the subcellular compartments (**Figure 4F**). We then plotted an average enrichment in axons versus soma for all annotated proteins in each overrepresented GO term in the category “cellular compartments”. This analysis showed that axonal proteome and transcriptome are associated with neuronal development and synaptic components, while somatic proteome and transcriptome – with the nuclear and endoplasmic reticulum (ER) components. Thus, our combined experiments demonstrate the feasibility of efficient separation of iMN-derived subcellular compartments for functional omics analyses.

### FUS^R244RR^-ALS hiPSC model recapitulates neurodegenerative phenotypes

To determine if iMNs can replicate the neurodegenerative phenotypes observed in ALS, we established a NIL-hiPSC line from the fibroblasts of an ALS patient with a mutation *FUS* gene (NIL-FUS^R244RR^-hiPSC). FUS is an RNA-binding protein playing a role in splicing and nucleo-cytosolic RNA transport and translation (reviewed in Piol et al., 2023). FUS^R244RR^ has a mutation in the RGG domain (**Figure 5A**), and mutations in this position have been reported to accelerate the conversion of FUS to a fibrous state, an event linked with disease (Patel et al., 2015). As a control, we created an isogenic line with the corrected FUS mutation (NIL-iso-hiPSC). We then differentiated these lines into iMNs and assessed their viability through a lactate dehydrogenase (LDH) assay. LDH is a cytosolic enzyme released into the cell culture medium upon loss of membrane integrity. LDH levels in the medium can be measured with a colorimetric assay. Our LDH measurements showed a ∼ 2.3-fold higher cell death of FUS^R244RR^-iMNs compared with the negative control isogenic iMNs (**Figure 5B**), pointing to the recapitulation of neurodegenerative phenotype in our test system.

**Figure 5.**
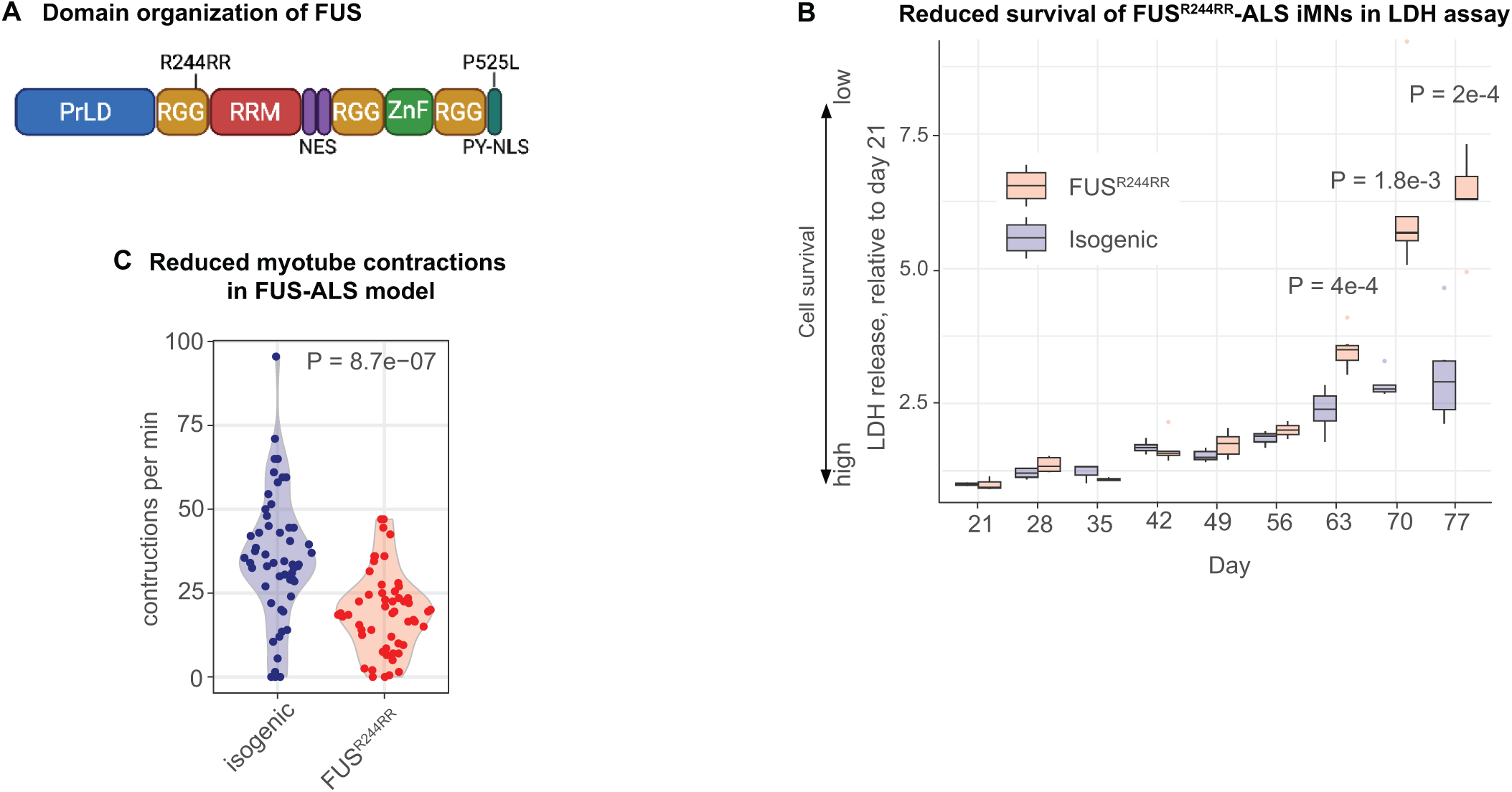
FUS^R244RR^-ALS hiPSC models recapitulate patient phenotypes. (**A**) Domain structure of FUS protein, including the positions of the examined R244RR mutation and the frequently researched P525L mutation (reviewed in Piol et al., 2023). PrLD: prion-like domain involved in FUS dimerization, RRG: arginine-glycine-glycine-rich domains, RRM: RNA recognition motif, NES: nuclear export signals, ZnF: zink-finger motif, PY-NLS: nuclear localization signal. RRM, ZnF, and RGG participate in RNA and protein binding. (B) The lactate dehydrogenase (LDH) release assay shows lower survival of FUS^R244RR^ iMNs (red), compared with isogenic control (blue). Boxplots showing LDH release, reflecting the level of plasma membrane damage (Y), at different time points (X). T-test P values for days 63, 70 and 77 are shown on the plot. **(C)** The number of myotube contractions per minute (X) is plotted for FUS^R244RR^-ALS (red) and control isogenic model (blue). P value was calculated using t-test.

ALS is characterised by progressive muscular paralysis reflecting degeneration of motor axons. To evaluate the progression of this degeneration in our ALS model, we used neuromuscular cultures. For that, we co-cultured either FUS^R244RR^-ALS iMNs or isogenic control iMNs with muscle cells derived from hiPSCs through the inducible expression of MyoD (Davis et al., 1987). We then counted the number of contractions in each co-culture. Notably, the FUS^R244RR^-ALS cultures exhibited a significantly reduced number of contractions compared to the isogenic controls, mirroring the impairment observed in patients (**Figure 5C**, **Table S4**, **Supplementary movies 2**). This analysis confirms that our neuromuscular cultures successfully recapitulate patient-specific phenotypes and can be effectively used to study the molecular mechanisms behind observed neurodegeneration defects.

### Spatial omics of hiPSC-derived ALS models

Next, we analyzed the changes in the local proteome of FUS^R244RR^-iMNs that could explain the reduced survival and functionality of mutant iMNs. We performed GO enrichment analysis of proteins differentially expressed in FUS^R244RR^ axons compared with isogenic axons (**Table S5**). Curiously, we detected axonal downregulation of proteins linked with neuromuscular junction (NMJ), extracellular matrix (ECM), soluble N-ethylmaleimide-sensitive factor activating protein receptor (SNARE) complex and axon guidance (**Figure 6A**, see **Table S6** for more details). These components are crucial for synapse and NMJ structure and plasticity (reviewed in Ferrer-Ferrer and Dityatev, 2018), and for simplicity, we refer to these proteins as axonal hits. The axonal hits include, for example, the Ephrin receptor EPHA4, a component of NMJ (Lai et al., 2001) whose knockout causes motor deficits in mice (Dominguez et al., 2020); the choline transporter SLC5A7, mutated in hereditary motor neuropathies (Salter et al., 2018); laminin A (LAMA1) whose neuronal knockdown results in abnormal morphology and neurite formation (Vilboux et al., 2016); laminin B (LAMB1) whose mutation in mice leads to dystonia-like movement disorders with spinal defects (Liu et al., 2015); and heparan sulfate proteoglycan 2 (HSPG2), which is aberrantly spliced in motor neurons of ALS patients (Rabin et al., 2010) and associated with tardive dyskinesia (Syu et al., 2010) (**Figure 6B**). Additionally, SNARE proteins, downregulated in FUS^R244RR^ axons, are crucial for the docking and fusion of synaptic vesicles with the plasma membrane (reviewed in Sauvola and Littleton, 2021) and thus are important for signal transmission to muscle cells (**Figure 6B**). These changes in the axonal proteome offer a mechanistic explanation for the observed functional defects in neuromuscular function, as detected in our contractility assay (**Figure 5C**).

**Figure 6.**
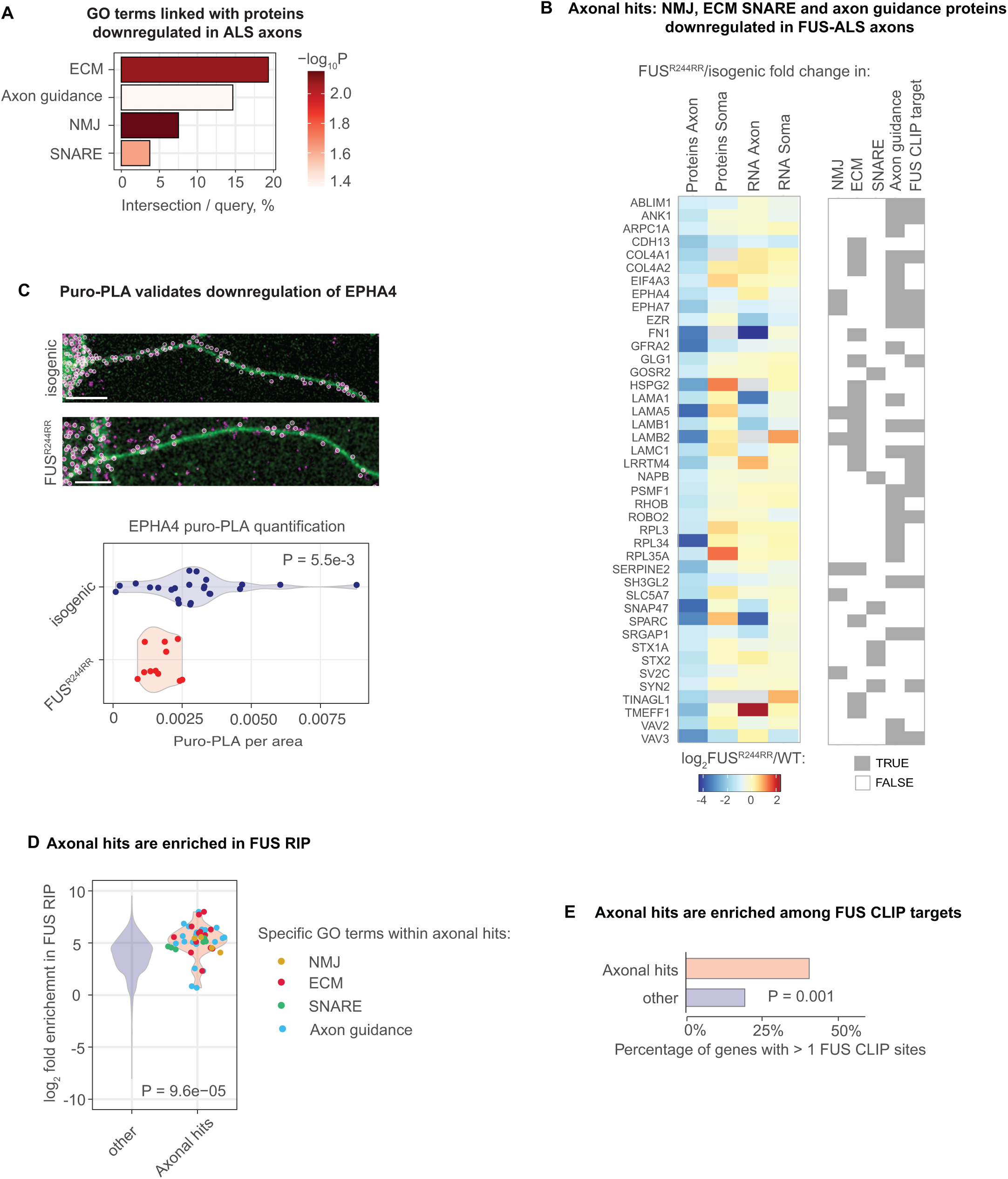
Proteins downregulated in FUS^R244RR^-ALS axons are linked in NMJ functionality and enriched in FUS targets. (A) GO terms associated with proteins downregulated in axons of FUS^R244RR^-ALS vs. isogenic iMNs. NMJ (neuromuscular junction), ECM (exracellular matrix), SNARE complex (soluble N-ethylmaleimide-sensitive factor activating protein receptor), involved in presynaptic membrane trafficking. **(B)** Heatmap showing changes in expression of proteins and RNAs belonging to downregulated GO term presented in (**A**) between FUS^R244RR^-ALS and isogenic iMNs (log2 fold change). The protein and RNA expression levels were measured by mass spectrometry and mRNA-seq analyses of axons and cell bodies. Specific GOs for each gene are indicated below the heatmap. This group of genes is further refered to as axonal hits. **(C)** Validation of EPHA4 downregualtion in FUS^R244RR^-ALS vs isogenic iMNs with puro-PLA. Representative EPHA4-puro-PLA images of FUS^R244RR^-ALS and isogenic iMN axons (above) and quantification of puro-PLA signal (below). EPHA4-puro-PLA: magenta; NF (serving to outline cell borders): green, scale bar: 10 µm. Circled are EPHA4-puro-PLA spots used for quantification. Quantification: Violin plots showing EPHA4-puro-PLA signal normalized per area in axons of FUS^R244RR^-ALS (red) vs isogenic (blue) iMNs. Individual datapoints correspond to single neurons (n(FUS^R244RR^-ALS) = 11, n(isogenic) = 24). P-value was calculated using t-test. **(D)** Axonal hits are enriched among FUS RIP targets. Violin plots showing fold enrichment (log2) of axonal hits (red) and all other RNAs (blue) in FUS RIP. FUS RIP data from (Reber et al., 2016) were used. Enrichment for indivudual RNAs within axonal hits is shown with a scatterplot and specific GO terms are indicated with color (NMJ: beige, ECM: red, SNARE: green, axon guidance: blue). **(E)** Axonal hits are enriched among FUS CLIP targets. Barplot showing percentage of FUS CLIP targets (> 1 CLIP site) among axonal hits (red) and other trasncripts (blue). FUS CLIP data are taken from Hoell et al. (2011).

Curiously, most axonal hits showed downregulation only in axons or stronger downregulation in axons than in cell bodies (**Figure 6B**). Such changes thus could not be detected with an analysis of whole MNs, so this experiment provides a proof of principle for the importance of subcellular analysis in ALS.

To understand if this downregulation is restricted to proteins or affects both proteins and mRNAs encoding them, we performed RNA-seq analysis of isolated subcellular compartments (**Table S7**). mRNAs of 30% of axonal hits (13 out of 42) showed downregulation in ALS motor axons; similarly to proteins, these transcripts were downregulated only in axons or more downregulated in axons than in cell bodies (**Figure 6B**). For the remaining 70%, the changes at the protein level could not be explained by changes in the levels of mRNAs encoding them (**Figure 6B**), suggesting that translation or protein localization may be affected. For example, NMJ component EPHA4 (Lai et al., 2001) is downregulated in FUS^R244RR^-ALS axons at the level of protein by ∼ 2-fold, but its mRNA levels are not significantly affected (**Figure 6B**). We therefore decided to test if the translation of EPHA4 is affected in FUS^R244RR^-ALS axons. We visualized its translation by the puro-PLA assay (tom Dieck et al., 2015), which relies on puromycin-tagging of newly synthesized proteins. Our results demonstrated that EPHA4 translation is indeed downregulated in FUS^R244RR^ axons compared with the isogenic control (**Figure 6C**).

Next, we sought to understand the mechanism behind the downregulation of these proteins in FUS^R244RR^ axons. FUS is an RNA-binding protein (RBP) that plays a role in multiple steps of RNA metabolism, including splicing, nucleo-cytosolic RNA transport, and translation (reviewed in Piol et al., 2023 7740). To explore if FUS may downregulate axonal hits by binding directly to transcripts encoding them, we analyzed data of FUS RNA-immunoprecipitation (RIP) from human cells (Reber et al., 2016) **Table S8**). Our analysis revealed that axonal hits, presented in **Figure 6B**, are enriched in FUS RIP (**Figure 6D**). CLIP (crosslinking immunoprecipitation) offers a significant advantage over RIP in that it enables the detection of RBP binding sites on RNA, thus allowing for the identification of direct RNA targets. We therefore also utilized FUS CLIP data (Hoell et al., 2011), **Table S9**). The Chi-squared test revealed a statistically significant enrichment of FUS CLIP sites among axonal hits (**Figure 6E**). Specifically, 40% of axonal hits contained more than one FUS CLIP site (17 out of 42), which was a 2-fold enrichment over other transcripts, in which 19% had more than one FUS CLIP site. The FUS RIP and CLIP data thus suggest that a substantial part of downregulated genes may represent direct FUS targets.

## DISCUSSION

As the global population ages, neurodegenerative diseases are projected by the World Health Organization to become the second leading cause of death by 2040, surpassing cancer-related deaths (reviewed in Gitler et al., 2017). Current therapeutic strategies primarily target specific genetic mutations responsible for these diseases. For example, antisense oligonucleotides (ASOs) are used to target and degrade aberrant mRNAs associated with various neurodegenerative diseases, such as ALS (targeting SOD1 and C9ORF72) (reviewed in Bennett et al., 2019; Gao et al., 2024). However, this approach has limitations due to the heterogeneous etiology of many neurodegenerative diseases. ALS, for instance, is linked to mutations in over 40 different loci, with SOD1 and C9ORF72 mutations accounting for only about 10% of all ALS cases (reviewed in Mathis et al., 2024). Moreover, for the majority of ALS patients, no traceable genetic cause can be identified, making targeted ASO therapies unsuitable for most cases. This highlights the necessity of understanding common defects across ALS cases of different etiologies.

Neurons are highly polarized cells with unique morphologies, comprising a cell body and neurite extensions—axons and dendrites. Axons play critical roles in conducting electrical signals and can be very long. Due to this specialized structure, many cellular components, including RNAs, proteins, and organelles, need to be transported into axons, where transported mRNAs are translated to produce local proteins (reviewed in Chekulaeva, 2024). This distinctive morphology is not found in any other cell type in the body and contributes to the vulnerability of neurons to neurodegenerative diseases. Compelling evidence shows that axonal loss is an early and predominant feature in multiple neurodegenerative diseases, preceding clinical symptoms and occurring before cell body degeneration (reviewed in Salvadores et al., 2017). This early event in the pathological process offers a potential therapeutic target.

Here, we established a new hiPSC-based model for ALS to study the mechanisms behind axonal dysfunction. We generated a hiPSC line from the fibroblasts of an ALS patient with an R2444RR mutation in the RGG domain of the *FUS* gene (FUS^R244RR^, **Figure 5A**). This is the first FUS-ALS model featuring this mutation. Although not identical, a mutation at the same position (R244C) has been studied in vitro and reported to accelerate the conversion of FUS droplets to a fibrous state (Patel et al., 2015). This model is crucial because most prior studies focused on FUS mutations affecting the NLS, causing protein mislocalization to the cytoplasm and leading to early disease onset. A diverse array of models is essential for a more comprehensive understanding of the disease pathogenesis.

We demonstrated that this model recapitulates phenotypes typical of ALS patients, such as reduced cell viability and reduced neuromuscular contractility (**Figure 5B-C**). Additionally, we developed an approach to efficiently isolate axonal and somatodendritic compartments from hiPSC-derived MNs for omics analysis. This analysis revealed a preferential downregulation of proteins linked with NMJ, ECM, SNARE complex, and axon guidance in FUS-ALS axons, which we termed axonal hits (**Figure 6A-B**). The axonal hits include, for example, the Ephrin receptor EPHA4, which mediates the phosphorylation of the actin-binding protein cortactin, thereby playing a role in NMJ maintenance (Lai et al., 2001). Consistently, EPHA4 knockout causes motor deficits in mice (Dominguez et al., 2020). Another hit is the choline transporter SLC5A7, which mediates choline uptake, a precursor of the MN neurotransmitter acetylcholine, and is mutated in hereditary motor neuropathies (Salter et al., 2018).

Crucially, these changes could not have been detected if the analysis had been performed on whole MNs, highlighting the importance of a subcellular analysis. Given the crucial role of these components in synapse and NMJ structure and plasticity, changes in the axonal proteome provide a mechanistic explanation for the observed neuromuscular functional defects detected in our contractility assay (**Figure 5C**). The exact mechanism by which mutant FUS leads to this dysregulation remains a key question for future studies.

Analysis of FUS RIP and CLIP data revealed that the FUS targets are enriched among the axonal hits, suggesting that many of these transcripts are directly regulated by FUS. Considering the involvement of FUS in RNA splicing (reviewed in Piol et al., 2023), we investigated whether the splicing of these transcripts is altered in FUS^R244RR^ iMNs. However, we found no splicing changes that could account for the altered expression of axonal hits in FUS^R244RR^-ALS (**Table S10**). Most axonal hits are affected at the protein level but not at the RNA level (**Figure 6B**), suggesting they are translationally downregulated in FUS^R244RR^-ALS axons. FUS is a multifunctional RBP known to influence multiple levels of gene expression, including translation (reviewed in Piol et al., 2023).

For instance, FUS has been reported to promote the translation of mRNAs localized to cellular protrusions in mouse fibroblasts with the assistance of the adenomatous polyposis coli (APC) protein (Yasuda et al., 2013). Conversely, ALS-linked FUS mutants were found to repress translation by forming heterogeneous condensates with fragile X mental retardation protein (FMRP) and sequestering bound RNAs from the translational machinery (Birsa et al., 2021). Similarly, another study reported that mutant FUS sequesters proteins associated with translation and RNA quality surveillance pathways, thereby suppressing protein translation (Kamelgarn et al., 2018). Additionally, ALS-linked FUS mutants have been shown to associate with stalled polyribosomes and reduce global protein synthesis (Sevigny et al., 2020). Activation of the integrated stress response (ISR) has also been reported in FUS-ALS neurons (Szewczyk et al., 2023). ISR is a key regulatory mechanism activated by various stress conditions (reviewed in Pakos-Zebrucka et al., 2016). It is triggered by the phosphorylation of the alpha subunit of eukaryotic initiation factor 2 (eIF2α), which in turn reduces the levels of the ternary complex eIF2:GTP:Met-tRNA^i^ required for translation initiation, leading to global translational inhibition. Notably, we have not detected an increase in eIF2α phosphorylation in our FUS-ALS model (data not shown). However, our model represents a mild form of ALS, with a mutation in the RGG domain rather than a mutation in the NLS domain, which results in early-onset ALS and has been the focus of the prior studies cited above.

In summary, we describe a novel mutation and establish a new patient-derived iPSC ALS model, demonstrate its impact on axonal gene expression, and provide a molecular explanation for the observed phenotype. Applying this methodology to other neurodegenerative models could elucidate shared pathways and molecular targets, paving the way for the development of broad-spectrum therapies aimed at restoring cellular functions and halting disease progression. This could result in more effective and accessible treatments, reducing the reliance on personalized therapies.

## METHODS

### Cell line generation

The stem cell line (Hues-3 HB9-GFP) was obtained from Harvard cell line collection (Di Giorgio et al., 2008). WT hiPSC line (BIH005-A) was obtained from the BIH-MDC stem core facility (https://hpscreg.eu/cell-line/BIHi005-A). ALS patient fibroblasts were collected at the ALS Clinic (Palermo) and reprogrammed into hiPSC by Applied Stem Cell and edited into wild type by Axol. To generate cell lines for differentiation into iMNs, the construct with the inducible NIL cassette (Fernandopulle et al., 2018) (Addgene #105841) was transfected into these lines using Lipofectamin-3000 and stably inserted into using TALENs (Addgene #62197 and #62196). To generate the cell line for differentiation into muscle cells (myoD-hiPSC), the construct (Addgene #105841) was modified to replace the NIL cassette with myoD coding sequence and stably integrated into BIH005-A using the same procedure.

### Cell culture and iMN differentiation protocol

hiPSC lines and Hues-3 line were grown in E8 medium, which was exchanged daily, and passaged before they reached full confluency. Differentiation was performed largely as described previously (Fernandopulle et al., 2018). For differentiation, cells were split with accutase from ∼80% confluent culture into Geltrex (Thermo A1413302) coated 6-well plates with 1-1.5 million cells per well in E8 with 10 μM ROCK inhibitor Y-27632 (Tocris 1254). At day 1 of differentiation medium was exchanged to induction medium (IM: DMEM/F12 w/ HEPES (Thermo 11330-032), N2 supplement (100x, Thermo 17502-048), Non-Essential Amino Acid (100x, Thermo 1140050), Glutamax (100x, Thermo 35050-061), ROCK inhibitor (10uM), Doxycycline (2 μg/ml), Compound E (0.1 μM, Enzo Life ALX-270-415-C250). On day 3 MN progenitors were replated using accutase, and the medium was exchanged to IM supplemented with 5-Fluoro-2’-deoxyuridine (FUDR, 8 μM, Sigma 50-91-9). Replating was done in accordance to the planned experiments: to Geltex-coated 6-well plates for isolation of RNA or protein from total cells, and to Geltrex-coated filter membrane inserts (pore size 1.0 μm PET membrane for 6-well plate Millicell PLRP06H48, Millipore) for isolation of RNA or protein from subcellular compartments or imaging on filters, to acid-treated and poly-D-lysine (100 μg/ml) and laminin (5 µg/ml) coated glass slides for immunofluorescence and imaging, or to poly-D-lysine (100 μg/ml) and laminin (5 µg/ml) coated CytoView MEA 48-plates (AXION Biosystems, M768-tMEA-48W) for direct electrophysiology measurements. On day 4 medium was exchanged to motor neuron media (MM: Neurobasal medium (Thermo 21103-049), B27 (50x, Thermo 17504-044), N2 (100x, Thermo 17502-048), Non-Essential Amino Acid (100x, Thermo 1140050), Glutamax (100x, Thermo 35050-061), Laminin (1 µg/ml), 10 ng/ml of BDNF (Peprotech 450-02), CNTF (Peprotech 450-13), and GDNF (Peprotech 450-10)), supplemented with doxycycline (1 µg/ml), Compound E (0.1 µM, Enzo Life ALX-270-415-C250) and FuDR (8 μM, Sigma 50-91-9). From then on, MM was exchanged every 2-3 days, and FUDR, Compound E and doxycycline were added to MM only until day 9.

### Harvesting of RNA and protein material

Protein material was harvested from filter membranes using 8 M UREA, 0.1 M Tris-HCl pH 7.5, and RNA material -using TRIfast (PeqLab 30-2010). For a detailed description of harvesting material from filter membranes see our prior work (Ludwik et al., 2019).

### hiPSC-derived neuromuscular culture

Myogenic differentiation was performed using published protocols (Sato et al., 2016; Shoji et al., 2016; Tanaka et al., 2013; Uchimura et al., 2017) with some modifications. Specifically, the myoD-hiPSC line, which contains myoD under a doxycycline-inducible promoter, was split using accutase. One million cells were plated per well of a Geltrex-coated 6-well plate in E6 medium (Thermo A1516401) supplemented with 10 μM ROCK inhibitor Y-27632 (Tocris 1254).

The next day (day 2), the medium was changed to E6 with 1 μg/ml doxycycline. On day 3, the cells were split using accutase, and 40.000 cells were plated per well of a Geltrex-coated 96-well square glass bottom plate (ibidi 89627) in KSR/αMEM medium. This medium consisted of 5% KnockOut Serum Replacement (KSR, Thermo 10828028), 55 mM 2-mercaptoethanol, 1x penicillin/streptomycin (Nacalai Tesque 2625384), and minimum essential medium α (MEMα, Thermo 12561056), and was additionally supplemented with 10 μM ROCK inhibitor and 1 μg/ml doxycycline.

The next day, the medium was changed to KSR/αMEM with 1 μg/ml doxycycline. Medium changes continued every second day until day 7. On day 8, the medium was switched to HS/DMEM, which included horse serum (Sigma H1138), DMEM high glucose (Thermo 11960085), 1x GlutaMAX (Thermo 35050-061), 55 mM 2-mercaptoethanol, and 1x penicillin/streptomycin, 10 ng/ml IGF-1 (PeproTech 100-11), and was additioanlly supplemented with 1 μg/ml doxycycline. Medium changes were performed every second day.

On day 14, 25.000 MN progenitors, generated via direct programming as described above, were co-plated per well in MM, supplemented with 1 μg/ml doxycycline and 10 ng/ml IGF-1. The medium was exchanged twice a week, with doxycycline included until day 20. Contractions were recorded at 6 weeks of myoD culture.

### Electrophysiology measurements using axion MEA plate

For electrophysiology measurements of iMN at day 21, the Maestro Edge multi-electrode array (MEA) system (AXION Biosystems) with CytoView MEA 48-plates (AXION Biosystems, M768-tMEA-48W) was used, and raw data was processed using AxIS Navigator software (version 2.0.4). Individual voltage spike events were extracted from the raw waveform of an exemplary electrode were analysed using the Spike Detector processing tool with a detection threshold of 5.5xSTD and then averaged for plotting.

### Lactate Dehydorgenase (LDH) assay

The LDH assay was conducted using the CyQuant LDH cytotoxicity assay kit (Thermo Fisher, C20300) according to the manufacturer’s instructions, with some modifications. In brief, iMNs were seeded onto a Geltrex-coated 96-well plate at a density of 187,500 cells/cm² and cultured following the iMN differentiation protocol. From day 21 onwards, media was changed weekly following the LDH measurement. LDH released by plasma-membrane-damaged iMNs into the media was quantified spectrophotometrically by measuring absorbance at 490 nm, with 680 nm used as the reference wavelength. Measurements from wells containing only media on the same plate served as a reference value.

### Immunofluorescence

For immunostaining, cells were grown on coverslips or ibidi plates as described above. Before staining, cells were fixed with 4 % paraformaldehyde for 20 min and permeabilized with 75 % EtOH or 0.2 % Triton X-100 in phosphate-buffered saline (PBS) for 10 min. For NMJ staining, 0.3% Triton X-100 in PBS was applied for 20 minutes. After blocking with 10% bovine serum albumin (BSA) in PBS for 60 min, cells were probed with the respective primary antibodies in 3% BSA in PBS overnight at 4°C, washed with 3 % BSA in PBS, and incubated with secondary antibodies for 1 h before mounting them with ProLong Gold with DAPI (Cell Signaling). The following primary antibodies were used: α-beta-III-tubulin (1:200, Sigma-Aldrich T2200), α-MAP2 (1:1.000, SYSY 188004), α-HB9 (1:25, DSHB 81.5C10), α-GFP (1:500, Abcam ab183734), α-Neurofilament (1:5.000, Biolegend BLD-822601; 1:1.000, Biolegend SMI312), α-Synapsin (1:500, Merck AB1543), α-myosin heavy chain (MyHC, 1:20, DSHB A4.1025). To visualize neuromuscular junctions, BTX-488 (1:100, Invitrogen B13422) was used. Images were taken using a Leica TCS SP8 confocal microscope with a 40x or 63x objective.

### Puro-PLA and image analysis

Detection of newly synthesized proteins by puro-PLA was performed as described previously (tom Dieck et al., 2015). Briefly, iMNs were incubated with 1 mg/ml puromycin for 15 min, washed quickly in PBS and fixed with 4% PFA in PBS for 10 min at RT. Cells were washed twice in PBS and permeabilized with 0.2 % Triton X-100 in PBS for 10 min at RT. Puro-PLA was performed using Duolink reagent, antibodies α-EPHA4 (1:100, Invitrogen PA514578), α-puromycin (1:200, Kerafast 3RH11), rabbit PLA^plus^ and mouse PLA^minus^ probes according to manufacturer’s recommendations except that antibody dilution solution was replaced by 5 % BSA in PBS. iMNs were immunostained with α-NF (1:1.000) and mounted in Duolink in situ mounting medium. Images were acquired using a 40x oil objective on a Leica SP8 FALCON confocal microscope.

The image analysis pipeline is adapted from a previously published workflow (Loedige et al., 2023) with the following adaptations. Max projections of the anti-Neurophillament and Puro-PLA images were generated using a Fiji (ImageJ) macro (Schindelin et al., 2012). Masks for the neurites were created using LABKIT (Arzt et al., 2022) and were manually corrected to represent either 30 -70 μm of the main neurites or the soma. The puro-PLA spots were detected and quantified in 2D max projections of the images using the RS-FISH Fiji plugin (Bahry et al., 2022). The detections were subsequently filtered using the binary neurite and soma masks in Python. The puro-PLA detections were normalized by the area of the neurite masks or soma masks. The pipeline can be found at: https://github.com/LauraBreimann/ALS_puro-PLA

### Electrophysiology calcium imaging

For calcium imaging experiments day 20 iMNs plated on 35 mm glass bottom imaging dish were loaded with 5μM CalBryte520 AM (AAT Bioquest) and 0.02% Pluronic acid in 50% Tyrodes buffer (pH 7.4, 120 mM NaCl, 2.5 mM KCl, 10 mM HEPES, 10 mM Glucose, 2 mM CaCl2, 1 mM MgCl2, osmolarity adjusted with sucrose to motor neuron media) and 50% Motor Neuron media for 30 min at 37 °C. All media was exchanged to Tyrodes buffer immediately before imaging. To induce action potentials (AP), a custom-made electrical stimulation chamber containing 2 electrodes was placed inside the iMN culture dish. The dish was mounted on a Dragonfly spinning disk microscope (Andor, Oxford Instruments) and connected to a pulse generator (A-M Systems Model 2100). After 5 seconds of baseline recording, iMN were stimulated by applying five field pulses at 9 s intervals, 1 ms width, and 20 V amplitude, followed by imaging up to 2.5 min to detect spontaneous firing. Imaging was performed using (488 nm with an exposure time of 100 ms) illumination in a 10x objective, and intracellular calcium dynamics were recorded at a frequency of 10 Hz. The post-acquisition analysis was performed using custom Matlab scripts, which normalized changes in fluorescence to the pre-stimulus baseline fluorescence. The last AP was plotted for each ROI, and the average of individual ROIs was calculated.

### Western Blot

For the western blot, 2.5 µg protein were separated on a 10% SDS-PAGE and then transferred to a PVDF membrane. The following primary antibodies were used to probe the membrane: α-Neurofilament (1:10.000, Biolegend BLD-822601), α-Synapsin (1:500, Merck AB1543), α-Synaptophysin (1:250, Life Technologies PA11043), α-Chat (1:100, Merck AB144P), α-beta-Actin (1:4.000, Sigma A2228), α-GAP43 (1:1.000, Santa Cruz sc-10786), α-RNA-PolII (1:200.000, Biolegend MMS-128P), α-H3 (1:3.000, Thermo PA5-31954).

### qPCR

RNA from total cells or separated compartments was treated with RQ1 Dnase and reverse-transcribed using the Maxima first strand cDNA synthesis kit (Thermo Fisher). qPCR was performed using sensiFAST SYBR No ROX qPCR kit (Bioline) and a CFX96 Real-Time PCR system (Biorad). Expression level changes were calculated using ΔΔCt method with beta-Actin as a reference RNA for total cells and expression normalised to day 1 timepoint or with rRNA as a reference RNA for separated compartments. The following primers were used: Actb-fw (GAGCACAGAGCCTCGCCTTT), Actb-rev (ACATGCCGGAGCCGTTGTC), rRNA-fw (AAACGGCTACCACATCCAAG), rRNA-rev (CCTCCAATGGATCCTCGTTA), NANOG-fw (ATGCCTCACACGGAGACTGT), Snord15b-fw (CAGTGATGACACGATGACGA), Snord15b-rev (AGGACACTTCTGCCAAAGGA), Rab13 (Primerbank ID 34850075c1), Kif1c (Primerbank ID 291327508c2), Kif5a (Primerbank ID 45446748c1), Gap43 (Primerbank ID 194248055c1).

### Mass spectrometry sample preparation

Samples for proteomics were harvested from total cells (one 6-well per replicate for time points day 1, 2, 3 and 4 and two wells per replicate for time points day 7, 14, and 21) and separated compartments (two 6-well membrane filters per sample) as described above and lysed in 8 M urea lysis buffer and processed for in-solution protein digestion (Mertins et al., 2018). All samples were prepared in triplicates. Briefly, di-sulfite bridges were reduced by treatment with 5 mM DTT for 1 h at room temperature and subsequently alkylated with 5 mM iodoacetamide (1 h, room temperature in the dark). Then samples were diluted with 50 mM Tris to reach a final urea concentration of 2 M, pre-digested with LysC (Wako Chemicals) at a 1:50 w:w ratio for 2 h, then trypsin (Promega) was added at 1:50 w:w ratio and samples were digested overnight at room temperature. Proteolytic digestions were stopped by acidification with the addition of formic acid to a final concentration of 1%. Digests were acidified with formic acid (FA) and centrifuged (20.000 g, 15 min) to remove the precipitated urea. The resulting peptides were de-salted via stop-and-go extraction (Rappsilber et al., 2007). 2 disks of C18 (3M Empore) material were inserted in a 200 μl pipette tip, activated via washes with methanol followed by washes with 50% acetonitrile (ACN) and 0.1 % FA and only 1 % FA. Samples were loaded onto the Stage-tips, and the retained peptides were washed twice with 0.1 % trifluoroacetic acid (TFA), followed by a wash with 1 % FA. Finally, peptides were eluted from the C18 material with a solution of 50 % ACN / 0.1 % FA and resuspended in mobile phase A (0.1 % FA and 3 % ACN in water) prior to mass spectrometric analysis.

### Liquid chromatography coupled with tandem mass spectrometry

Approximately 1 μg of peptides for each sample was online-separated on an EASY-nLC 1200 (Thermo Fisher Scientific) and acquired on a Q-Exactive HFx (Thermo Fisher Scientific), and samples were run with a randomised order. Peptides were separated on a fused silica, 25 cm long column packed in-house with C18-AQ 1.9 μm beads (Dr. Maisch Reprosil Pur 120) kept at a temperature of 45 °C. After equilibrating the column with 5μl mobile phase A, peptides were separated with a 250 μl/min flow on a 214 min gradient: mobile phase B (0.1 % FA and 90 % ACN) increased from 2 % to 30 % in the first 192 min, followed by an increase to 60 % in the following 10 min, to then reach 90 % in one min, which was held for 5 min. The mass spectrometer was operated in data-dependent acquisition, with MS1 scans from 350 to 1500 m/z acquired at a resolution of 60,000 (measured at 200 m/z), maximum injection time (IT) of 10 ms, automatic gain control (AGC) target value of 3 × 10^6^ and recording in profile mode. The 20 most intense precursor ion peaks with charges from +2 to +6 were selected for fragmentation, unless present in the dynamic exclusion list (30 s). Precursor ions were selected with an isolation window of 1.3 m/z, fragmented in an HCD cell with a normalised collision energy of 26 %, and analysed in the detector with a resolution of 15.000 m/z (measured at 200 m/z), AGC target value of 10^5^, maximum IT of 22 ms and recording in centroid mode.

### Proteomics data analysis

RAW files were analysed using MaxQuant (Tyanova et al., 2016) version 1.6.3.4 using MaxLFQ as a quantification method (Cox et al., 2014). The MS scans were searched against the human Uniprot databases (Jan 2020) using the Andromeda search engine with FDR calculated based on searches on a pseudo-reverse database and set to 0.05. The search included as fixed modifications carbamidomethylation of cysteine and as variable modifications methionine oxidation, N-terminal acetylation, and asparagine and glutamine deamidation. Trypsin/P was set as a protease for in-silico digestion. All samples (time points and soma/neurite comparisons) were analysed in the same MaxQuant session. The resulting protein groups were filtered for protein contaminants, hits in the reverse database, only identified by modified site and identified by less than two peptides, of which one was unique. In the analysis of the time course experiment, we also filtered out proteins that were not identified in all replicates in at least one time point. The remaining missing values were imputed by randomly selecting values from a normal distribution with a 30% the standard deviation and shifted downwards by 1.8 standard deviation units (Hein et al., 2015).

For analysis of expression levels imputed LFQ values were log-transformed, averaged between replicates and protein identifiers mapped to ensembl gene IDs. For heatmaps values were also scaled to z-scores (number of standard deviations from the mean) for each protein across all time points.

In the analysis of the cellular compartments, proteins were selected when present in all replicates of one cellular compartment, and the remaining missing values were imputed as described above. Proteins differentially expressed between soma and neurite were evaluated with a modified Student t-test (Hein et al., 2015) with an S0 value of 1. Statistical analysis was done with R (v3.6.3). P-values were set to 1 for proteins detected in less than two samples of any compartment.

### RNA-seq libraries

100 ng of total RNA obtained from either total cells or separated compartments was used for library preparation with the Truseq stranded total library prep kit (Illumina 20020596) or Truseq stranded mRNA library prep kit (Illumina 20020595) according to the manufacturer’s recommendation. Each library was prepared in triplicate and sequenced on an Illumina NextSeq 500 sequencer with single-end 150 bp reads.

### RNA-seq data analysis

All fastq files were trimmed, mapped & counted using the PiGx RNAseq pipeline (Wurmus et al., 2018) version 0.0.10). For analysis of expression values raw counts generated by salmon were scaled to transcripts-per-million (TPM) and genes with average TPM values below one were removed before log-transformation of TPM values. Some of our individual samples that did not cluster with replicates in PCA were removed before any further analysis. For heatmaps, TPM values were also scaled to z-scores (number of standard deviations from the mean) for each gene across all timepoints. Differential expression between compartments was performed using the PiGx pipeline, but only genes with TPM > 1 in at least two replicates were retained for analysis.

For alternative splicing analysis, STAR alignment of the RNA-seq data was fed through the LeafCutter pipeline (Li et al., 2018) with default parameters. Events with P < 0.05 were considered as significant.

Public available datasets were obtained from the following accessions: GSE108094 (control samples SRR6376956-59, Rizzo et al., 2019), GSE69175 (control samples SRR2038215-16, Ng et al., 2015), GSE121069 (human control SRR7993125-37, Nijssen et al., 2018), SRP064478 (healthy controls SRR2558717-24, Brohawn et al., 2016), GSE41795 (positive samples SRR606335-36, Amoroso et al., 2013), GSE66230 (SRR1814073,75-76,78,79-80, Briese et al., 2016). Fastq files were processed using the same PiGx workflow as before. Raw count values were extracted from supplementary files of Maciel et al. (2018) and Rotem et al. (2017). Variance stabilizing transformation was applied to raw counts from all datasets for those genes with per-dataset average TPM values above 1 using DESeq2. PCA was then performed on median values from the total or soma compartments of each dataset.

### Gene ontology analysis

GO analysis was performed using the gprofiler2 R package (Kolberg et al., 2020) call to web-interface, 06/2020) on genes that had significant enrichment (p.adj < 0.05) with |log2fc| > 1 in either soma or axon compartment on both RNA and protein (MS) level. GO terms enriched in the localised gene set from both compartments were then filtered for a maximum of 1000 genes per term and at least 25 overlapping localised genes/proteins with each term and further based on the graph of GO term relationships: for each connected sub-tree of enriched terms (nodes), only the lowermost (no enriched direct daughter terms) and uppermost (no direct enrich parent terms) were retained (GO release 2020-06). For each such selected GO term protein localisation values (log2fc) from all genes annotated as members of this term were used for the generation of figures.

### Cell type deconvolution

Raw fastq files for single cells as well as their AP type classification were provided by Bardy et al. (Bardy et al., 2016). Files were processed using PiGx RNAseq pipeline, and raw counts from salmon were imported using tximport. Seurat was used to process (CreateSeuratObject, parameters: min.cells = 3) and normalise counts, and only cells with from AP type 1-5 were used, and types 1-3 were grouped together. Differentially expressed marker genes between AP type groups (1-2-3, 4 & 5) were then calculated using FindAllMarkers (parameters: logfc.threshold = 0.25 and min.pct = 0.25).

All markers identified this way were used as signature markers for cell type deconvolution of total cell time points day 4 -day 21 with CIBERSORTx (Non-default signature matrix settings: 25-300 barcode genes; single cell min. expression 1, replicates 0 and sampling 0; 500 permutations used for statistical analysis in cell fraction imputation, (Newman et al., 2019).

## Data availability

The mass spectrometry proteomics data have been deposited to the ProteomeXchange Consortium via the PRIDE partner repository with the dataset identifier PXD031066 and PXD054876. Raw RNA-seq data have been deposited to EBI ArrayExpress with identifiers E-MTAB-14352, E-MTAB-14354, and E-MTAB-14359. The pipeline for puro-PLA data analysis can be found at: https://github.com/LauraBreimann/ALS_puro-PLA. Supplementary movies are deposited at Figshare with doi 10.6084/m9.figshare.26412352 (Supplementary movie 1) and 10.6084/m9.figshare.26503363 (Supplementary movies 2).

## ACKNOWLEDGMENTS

We thank the MDC microscopy and stem core facilities for technical support. This work was supported by the EU-JPND grant to M.C., V.B. and I.U.

## AUTHOR CONTRIBUTIONS

The experiments were performed by K.L. (generation of NIL-hiPSC and NIL-hESC cell lines; preparation of RNA-seq libraries and samples for proteomics analysis in Fig 2A-B, 4, S1, S2; western blots in Fig 2C, 4C; rt-qPCR in Fig 4E; IF and imaging in Fig 1B, 1E, 4B), S.M. (preparation of RNA-seq libraries and samples for proteomics analysis for FUS-ALS model in Fig 6B, IF 1C, calcium imaging in Fig 3B-C), C.R. (LDH assays in Fig 5B; IF in Fig 1D; puro-PLA image acquisition in Fig 6C), E.B. (generation of myoD-hiPSC line, electrophysiology in Fig 3D), M.S. (neuromuscular culture and quantification of contraction in Fig 5E), T.M. and M.S. (mass spetrometry), S.M. (NMJ imaging in Fig 3E). V.B. provided patient fibroblasts. E.R, K.L and C.R. contributed to insertion of NIL cassete into FUS^R244RR^ hiPSC and isogenic lines. N.G. contributed to testing protocols for neuromuscular cultures and reported the differences in contractility of mutant and control cultures. Analysis was performed by N.K. (data analysis and plotting in Fig 2A-B, 2D, 3A, 4F, 6A, S1, S2), I.U. (data analysis for FUS RIP in Fig 6D, GO terms, splicing), M.C. (exploratory analysis in Fig 4D, 6B, 6E, plotting in Fig 5B-C, 6C-D) and L.B. (puro-PLA quantification in Fig 6C). A.O.M. developed the protocol for calcium imaging, and A.W. performed data analysis (in Fig 3B-C). Figures were assembled by N.K., K.L. and M.C. The manuscript was written by N.K. and M.C. with contributions from other co-authors.

## Competing financial interest

The authors declare no competing financial interest.

**Figure S1. Quality control plots for transcriptomic and proteomic data from iMN time course experiment.**

(A-B) Correlation plots of expression of RNA (A, log10(TPM), with 0.01 pseudocount) and protein (B, logLFQ) in different samples taken from iMN at different stages of differentiation. The number shown in each rectangle represents the Pearson correlation coefficient between the samples.

(C-D) The principle component analysis of iMN expression data from different time points. PCA was calculated using only genes/proteins detected in all samples based on (C) log10(TPM) and (D) logLFQ values.

**Figure S2. Quality control plots for transcriptomic and proteomic data of separated iMN compartments.**

(A-B) Correlation plots of expression of RNA (A, log10(TPM), with 0.01 pseudocount) and protein (B, logLFQ) in axonal and somatodendritic compartments from iMNs at day 21 of differentiation. The number shown in each rectangle represents the Pearson correlation coefficient between the samples.

(C-D) The principle component analysis of axon and soma compartment from iMN expression data at day 21 after differentiation. PCA was calculated using only genes/proteins detected in all samples based on (C) log10(TPM) and (D) logLFQ values.

(E-F) MA plots showing the distribution of log2 fold change values of proteins (E) and RNA (F) between axonal and somatodendritic compartments in relation to mean abundance (log2LFQ for proteins and log2TPM for RNA). Color indicates significant enrichment (p.adj < 0.05 & |log2fc| > 1) in either compartment (axons: purple; soma&dendrites: blue).

## List of Supplementary tables

Table S1. Mass spectormetry and RNA sequencing data for different differentiation stages of iMNs.

Table S2. Mass spectormetry data for axonal and somatodendritic compartments of iMNs.

Table S3. RNA-seq data for axonal and somatodendritic compartments of iMNs.

Table S4. Contractions in FUS(R244RR) and isogenic control in neuromuscular cultures.

Table S5. Mass spectormetry data for axonal and somatodendritic compartments of FUS(R244RR) and isogenic control iMNs.

Table S6. Gene Ontology analysis of differentially expressed axonal proteome in FUS(R244RR) versus isogenic control iMNs.

Table S7. RNA-seq data for axonal and somatodendritic compartments of FUS(R244RR) and isogenic control iMNs.

Table S8. FUS RNA Immunoprecipitation data ((Reber et al., 2016)). Table S9. FUS CLIP data (Hoell et al., 2011)).

Table S10. Differential splicing analysis in FUS(R244RR) and isogenic control iMNs.

