## Supplementary material for "Neuromuscular dysfunction in patient-derived FUS^R244RR^-ALS iPSC model via axonal downregulation of neuromuscular junction proteins": Figure S1

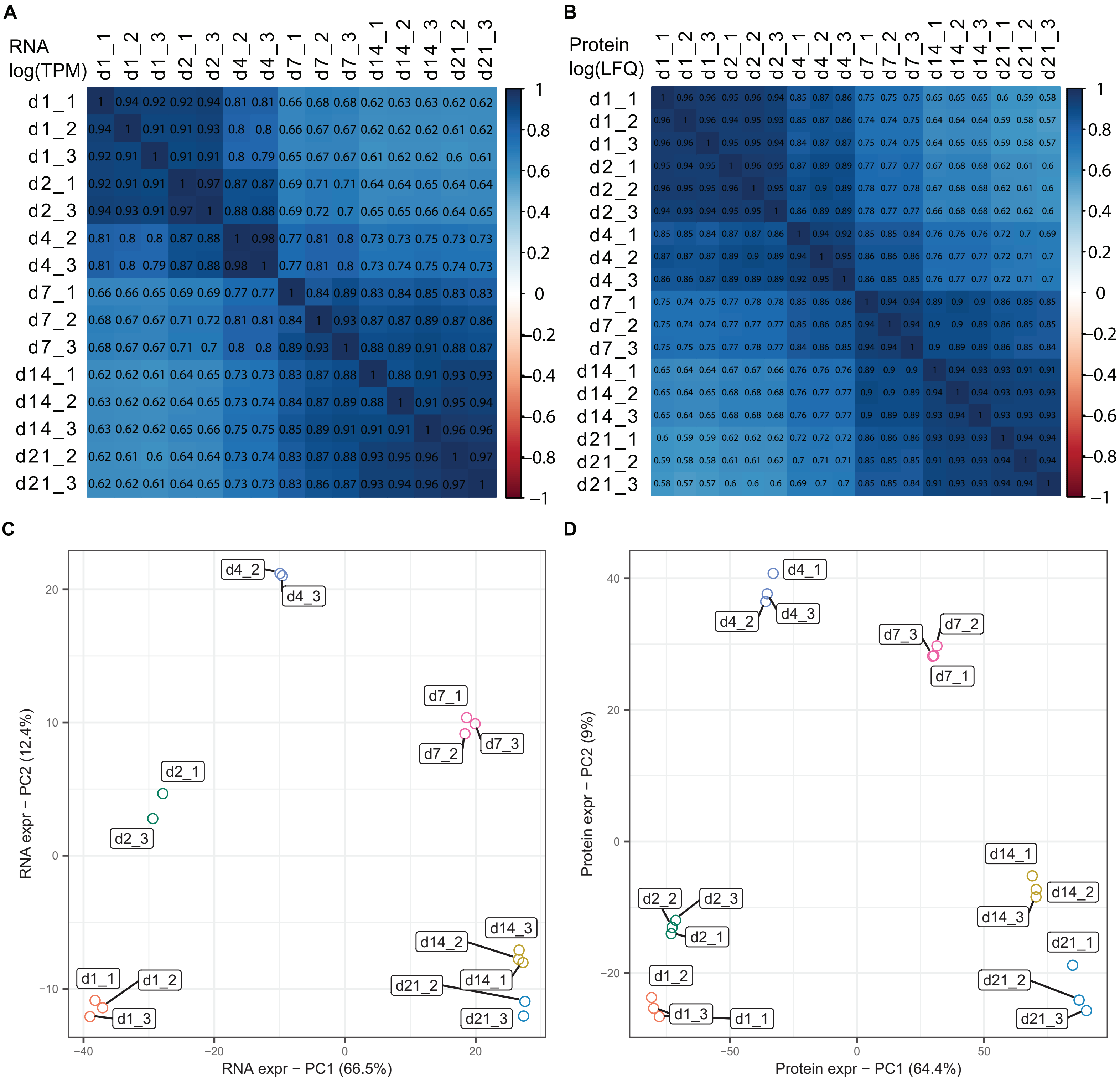


**Figure S1. Quality control plots for transcriptomic and proteomic data from iMN time course experiment.**
(A-B) Correlation plots of expression of RNA (A, log10(TPM), with 0.01 pseudocount) and protein (B, logLFQ) in different samples taken from iMN at different stages of differentiation. The number shown in each rectangle represents the Pearson correlation coefficient between the samples.
(C-D) The principle component analysis (PCA) of hiMN expression data from different timepoints. PCA was calulated using only genes/proteins detected in all samples based on (C) log10(TPM) and (D) logLFQ values.
