## Supplementary material for "Neuromuscular dysfunction in patient-derived FUS^R244RR^-ALS iPSC model via axonal downregulation of neuromuscular junction proteins": Figure S2

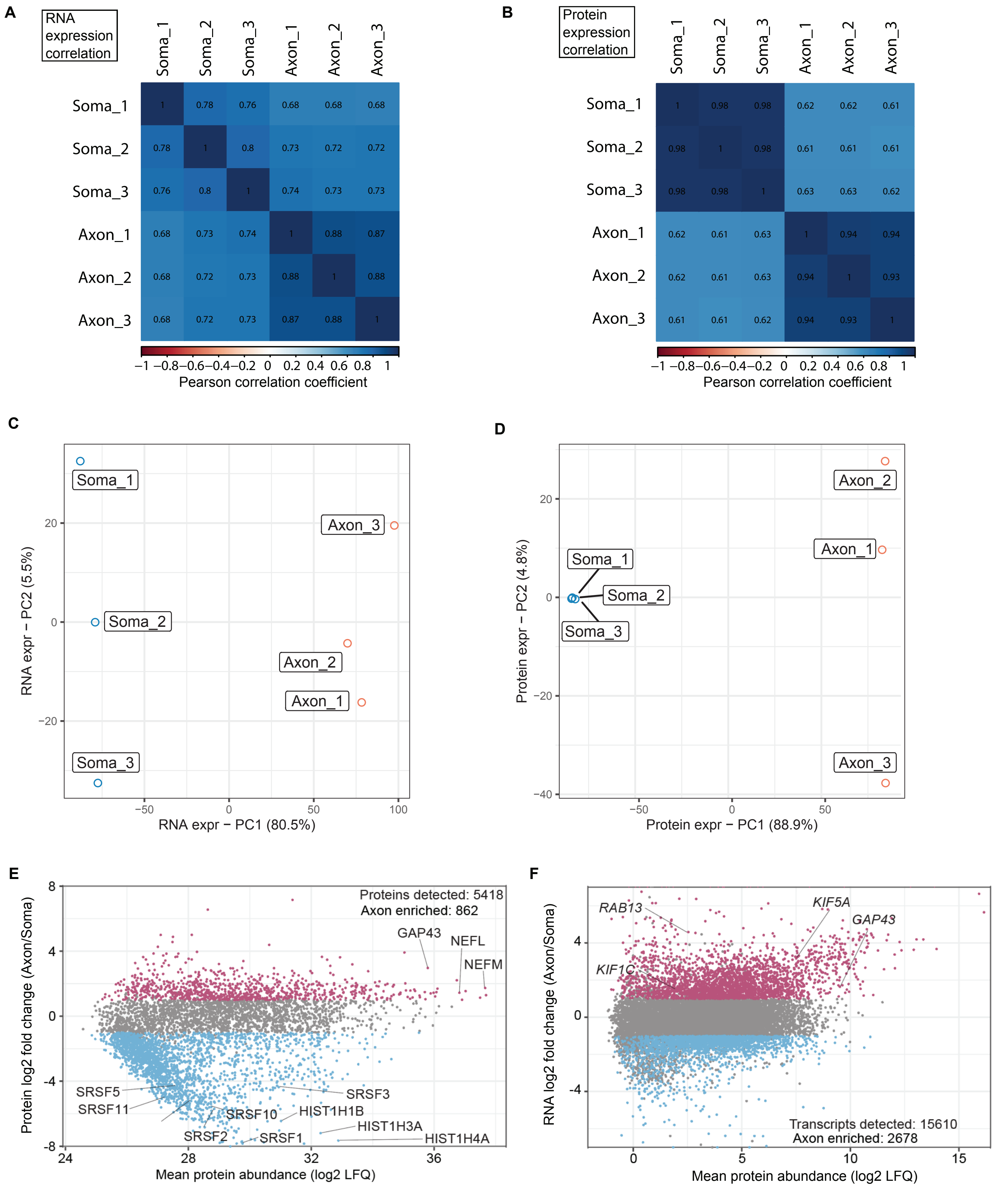


**Figure S2. Quality control plots for transcriptomic and proteomic data of separated iMN compartments.**

(A-B) Correlation plots of expression of RNA (A, log10(TPM), with 0.01 pseudocount) and protein (B, logLFQ) in axonal and somatodendritic compartments from iMNs at day 21 of differentiation. The number shown in each rectangle represents the Pearson correlation coefficient between the samples.

(C-D) The principle component analysis of axon and soma compartment from iMN expression data at day 21 after differentiation. PCA was calculated using only genes/proteins detected in all samples based on (C) log10(TPM) and (D) logLFQ values.

(E-F) MA plots showing the distribution of log2 fold change values of proteins (E) and RNA (F) between axonal and somatodendritic compartments in relation to mean abundance (log2LFQ for proteins and log2TPM for RNA). Color indicates significant enrichment ($p.adj < 0.05$ & $|log2fc| > 1$) in either compartment (axons: purple; soma&dendrites: blue).
